# Connections between freshwater carbon and nutrient cycles revealed through reconstructed population genomes

**DOI:** 10.1101/365627

**Authors:** Alexandra M. Linz, Shaomei He, Sarah L. R. Stevens, Karthik Anantharaman, Robin R. Rohwer, Rex R. Malmstrom, Stefan Bertilsson, Katherine D. McMahon

## Abstract

Metabolic processes at the microbial scale influence ecosystem functions because microbes are responsible for much of the carbon and nutrient cycling in freshwater. One approach to predict the metabolic capabilities of microbial communities is to search for functional marker genes in metagenomes. However, this approach does not provide context about co-occurrence with other metabolic traits within an organism or detailed taxonomy about those organisms. Here, we combine a functional marker gene analysis with metabolic pathway prediction of microbial population genomes (MAGs) assembled from metagenomic time series in eutrophic Lake Mendota and humic Trout Bog to identify how carbon and nutrient cycles are connected in freshwater. We found that phototrophy, carbon fixation, and nitrogen fixation pathways co-occurred in *Cyanobacteria* MAGs in Lake Mendota and in *Chlorobiales* MAGs in Trout Bog. *Cyanobacteria* MAGs also had strong temporal correlations to functional marker genes for nitrogen fixation in several years. Genes encoding steps in the nitrogen and sulfur cycles varied in abundance and taxonomy by lake, potentially reflecting the availability and composition of inorganic nutrients in these systems. We were also able to identify which populations contained the greatest density and diversity of genes encoding glycoside hydrolases. Populations with many glycoside hydrolases also encoded pathways for sugar degradation. By using both MAGs and marker genes, we were better able to link functions to specific taxonomic groups in our metagenomic time series, enabling a more detailed understanding of freshwater microbial carbon and nutrient cycling.

## Introduction

Lakes collect nutrients from surrounding terrestrial ecosystems (Williamson et al., 2008), placing lakes as “hotspots” for carbon and nutrient cycling in the landscape (Butman et al.,2015). Much of this biogeochemical cycling is performed by freshwater microbes. We have learned much about freshwater microbes through previous research that has revealed high levels of diversity and change over time in freshwater microbial communities (Allgaier & Grossart, 2006), the geographic distribution of freshwater taxa (Šimek et al., 2010), the distribution of functional marker genes (Peura et al., 2012, 2015; Ramachandran & Walsh, 2015; Eiler et al., 2016), and substrate use capabilities in specific phylogenetic groups (Salcher, Posch & Pernthaler, 2013). However, organism-level information about microbial metabolism is currently not well incorporated into conceptual models of freshwater carbon and nutrient cycling.

Although aquatic microbes are often classified either exclusively as decomposers or phytoplankton, their roles and relative importance in the food chain are now recognized as distinct and complex (Pomeroy & Wiebe, 1988). Dissolved organic carbon (DOC) is produced at every trophic level, but this carbon is often not in a form directly available for consumption by secondary or tertiary trophic levels. Instead, microbes are responsible for processing this complex, recalcitrant DOC, producing more labile biomass that is subsequently consumed. This process of maintaining DOC within the food web is known as the “microbial loop” (Azam et al., 1983), although aquatic microbes respire much of the DOC to CO_2_. In some systems, microbial respiration is thought to exceed primary production, resulting in the release of excess of CO2 to the atmosphere (del Giorgio, Cole & Cimbleris, 1997). Inorganic compounds can be used as nitrogen and sulfur sources, or they can provide energy to chemolithotrophs that are in turn consumed by other trophic levels. Microbial conversions of inorganic compounds are often just as crucial to freshwater biogeochemistry as the degradation of DOC.

Previously, we used time series metagenomics to assemble nearly 200 metagenome-assembled genomes (MAGs) from two temperate lakes: Lake Mendota, a highly productive eutrophic lake, and Trout Bog, a humic bog lake (Bendall et al., 2016). These MAGs were used to study genome-wide diversity sweeps in Trout Bog (Bendall et al., 2016), to build metabolic networks of the ubiquitous freshwater *Actinobacteria* acI (Hamilton et al., 2017), and to propose functions for freshwater *Verrucomicrobia* (He et al., 2017). In addition to this body of knowledge based on the MAG dataset, previous time series analyses of 16S rRNA gene amplicon datasets from both lakes provide an understanding of taxon dynamics over time (Hall et al., 2017; Linz et al., 2017). Lake Mendota and Trout Bog are ideal sites for comparative time series metagenomics because of their history of extensive environmental sampling by the North Temperate Lakes - Long Term Ecological Research program (NTL-LTER, http://lter.limnology.wisc.edu) and their contrasting limnological attributes (Table 1, Table S1). Here, we build on this previous work by identifying contrasting patterns of carbon and nutrient cycling between the lakes based on analyses of functional marker genes and MAGs.

Gene-centric methods are one method that can identify community functions, while analysis of population genomes using MAGs can identify coupled metabolic processes taking place within the boundary of a cell. In this research, we use functional marker genes and MAGs from two freshwater lakes with contrasting chemistry to yield insights about microbial metabolism in freshwater ecosystems. We identified genes and pathways purportedly involved in primary production, DOC mineralization, and nitrogen and sulfur cycling. Some types of metabolisms were found in both sites despite their different chemistry profiles, but in different taxonomic groups. We demonstrate how MAGs and metagenomic time series can be used to track specific phylogenetic groups capable of key biogeochemical transformations. Finally, we introduce the MAG collection as a valuable community resource for other freshwater microbial ecologists to mine and incorporate into comparative studies across lakes around the world.

## Methods

### Sampling

Samples were collected from Lake Mendota and Trout Bog as previously described (Bendall et al., 2016). Briefly, integrated samples of the water column were collected during the ice-free periods of 2007-2009 in Trout Bog and 2008-2012 in Lake Mendota. In Lake Mendota, the top 12 meters of the water column were sampled, approximating the epilimnion (upper, oxygenated, and warm thermal layer). The epilimnion and hypolimnion (bottom, anoxic, and cold thermal layer) of Trout Bog were sampled separately at depths determined by measuring temperature and dissolved oxygen concentrations. The sampling depths were most often 0-2 meters for the epilimnion and 2-7 meters for the hypolimnion. DNA was collected by filtering 150 mL of the integrated water samples on 0.2-μm pore size polyethersulfone Supor filters (Pall Corp., Port Washington, NY, USA). Filters were stored at −80C until extraction using the FastDNA Spin Kit (MP Biomedicals, Burlingame, CA, USA).

### Sequencing

As previously described (Bendall et al., 2016; Roux et al., 2017), metagenomic sequencing was performed by the Department of Energy Joint Genome Institute (DOE JGI) (Walnut Creek, CA, USA). A total of 94 samples were sequenced for Lake Mendota, while 47 metagenomes were sequenced for each layer in Trout Bog. Samples were sequenced on the Illumina HiSeq 2500 platform (Illumina, San Diego, CA, USA), except for four libraries (two from each layer of Trout Bog) that were sequenced using the Illumina TruSeq protocol on the Illumina GAIIx platform (Data S1). Paired-end sequencing reads were merged with FLASH v1.0.3 with a mismatch value of less than 0.25 and a minimum of 10 overlapping bases, resulting in merged read lengths of 150-290 bp (Magooc & Salzberg, 2011). 16S rRNA gene amplicon sequencing was also performed on samples collected with the same method over the same time periods. This data is available under DOE JGI project IDs 1078703 and 1018581 for Trout Bog and Lake Mendota, respectively. Samples from Trout Bog were sequenced on the 454 GS FLX-Titanium platform (Roche 454, Branford, CT, USA) targeting the V8 hypervariable region (primer 1392R: ACGGGCGGTGTGTRC) (Engelbrektson et al., 2010), and sequences were trimmed to 324 base pairs using VSEARCH (v2.3.4) (Rognes et al., 2016). Samples from Lake Mendota were sequenced on an Illumina MiSeq, and the V4 region was targeted using paired-end sequencing (primers 525F: GTGCCAGCMGCCGCGGTAA and 806R: GGACTACHVGGGTWTCTAAT) (Caporaso et al., 2012). Both datasets were trimmed based on alignment quality and chimera checking using mothur v.1.39.5 (Schloss et al., 2009). Unclustered unique sequences were assigned taxonomy using TaxAss (Rohwer et al., 2017) to leverage the FreshTrain (version FreshTrain25Jan2018Greengenes13_5) (Newton et al., 2011) and Greengenes (version 13_5) (DeSantis et al., 2006).

### Assembly and Binning

To recover MAGs, metagenomic reads were pooled by lake and layer and then assembled as previously described (Bendall et al., 2016; Roux et al., 2017). In Trout Bog, this assembly was performed using SOAPdenovo2 at various k-mer sizes (Luo et al., 2012), and the resulting contigs were combined using Minimus (Sommer et al., 2007). In Lake Mendota, merged reads were assembled using Ray v2.2.0 with a single k-mer size (Boisvert et al., 2012). Contigs from the combined assemblies were binned using MetaBAT (-veryspecific settings, minimum bin size of 20kb, and minimum contig size of 2.5kb) (Kang et al., 2015), and reads from unpooled metagenomes were mapped to the assembled contigs using the Burrows-Wheeler Aligner (≥ 95% sequence identity, n = 0.05) (Li & Durbin, 2010), which allowed time-series resolved binning (Table S2). DOE JGI’s Integrated Microbial Genome (IMG) database tool (https://img.jgi.doe.gov/mer/) (Markowitz et al., 2012) was used for gene prediction and annotation. Annotated MAGs can be retrieved directly from the IMG database and JGI’s Genome Portal using the IMG Genome ID provided (also known as IMG Taxon ID). MAG completeness and contamination/redundancy was estimated based on the presence of a core set of genes with CheckM (Rinke et al., 2013; Parks et al., 2015), and MAGs were classified using Phylosift (Darling et al., 2014) or the phylogeny-based “guilt by association” method (Hamilton et al., 2017).

### Functional Marker Gene Analysis

To analyze functional marker genes in the unassembled, unpooled metagenomes, we used a curated database of reference protein sequences (Data S2) (Anantharaman et al., 2016) and identified open reading frames (ORFs) in our unassembled metagenomic time series using Prodigal (Hyatt et al., 2010). This analysis was conducted on merged reads. The protein sequences and ORFs were compared using BLASTx (Camacho et al., 2009) with a cutoff of 30% identity. Significant differences in gene frequency between sites were identified using LEfSE (Segata et al., 2012). Read abundance was normalized by metagenome size for plotting. We chose to perform this analysis because gene content in unassembled metagenomes is likely more quantitative and more representative of the entire microbial community than gene content in the MAGs.

### Pathway Prediction

Only MAGs that were at least 50% complete with less than 10% estimated contamination (meeting the MIMARKS definition of a medium or high quality MAG) were included in this study (Bowers et al., 2017). Taxonomy was assigned to MAGs using Phylosift (Darling et al., 2014). Pathways were analyzed by exporting IMG’s functional annotations for the MAGs, including KEGG, COG, PFAM, and TIGRFAM annotations and mapped to pathways in the KEGG and MetaCyc databases as previously described (He et al., 2017). To score presence, a pathway needed at least 50% of the required enzymes encoded by genes in a MAG and if there were steps unique to a pathway, at least one gene encoding each unique step. Putative pathway presences was aggregated by lake and phylum in order to link potential functions identified in the metagenomes to taxonomic groups that may perform those functions in each lake. Glycoside hydrolases were annotated using dbCAN (http://csbl.bmb.uga.edu/dbCAN) (Yin et al., 2012). Nitrogen usage in amino acids was calculated by taking the average number of nitrogen atoms in translated ORF sequences across each MAG.

Data formatting and plotting was performed in R (R Core Team (2017). R: A language and environment for statistical computing. R Foundation for Statistical Computing, Vienna, Austria. URL https://www.R-project.org/.) using the following packages: ggplot2 (H. Wickham. ggplot2: Elegant Graphics for Data Analysis. Springer-Verlag New York, 2009.), cowplot (Claus O. Wilke (2017). cowplot: Streamlined Plot Theme and Plot Annotations for ‘ggplot2’. R package version 0.9.2. https://CRAN.R-project.org/package=cowplot), reshape2 (Hadley Wickham (2007). Reshaping Data with the reshape Package. Journal of Statistical Software, 21(12), 1-20. URL http://www.jstatsoft.org/v21/i12/.), and APE (Paradis E., Claude J. & Strimmer K. 2004. APE: analyses of phylogenetics and evolution in R language. Bioinformatics 20: 289-290.). The datasets, scripts, and intermediate files used to predict pathway presence and absence are available at <https://github.com/McMahonLab/MAGstravaganza>. Any future updates or refinements to this dataset will be available at this link.

## Results/Discussion

### Community Functional Marker Gene Analysis

To assess potential differences in microbial metabolisms between Lake Mendota and Trout Bog, we tested whether functional marker genes identified in the unassembled merged metagenomic reads appeared more frequently in one lake or layer compared to the others. These comparisons were run between the epilimnia of Trout Bog and Lake Mendota, and between the epilimnion and hypolimnion of Trout Bog. We did not compare the epilimnion of Lake Mendota to the hypolimnion of Trout Bog, as the multitude of factors differing between these two sites make this comparison illogical. Many genes differed significantly by site, indicating contrasting gene content between lakes and layers (Data S3). To further infer differences in microbial metabolism, we aggregated marker genes by function (as several marker genes from a phylogenetic range were included in the database for each type of function) and tested for significant differences in distribution between lakes and layers using a Wilcoxon rank sum test with a Bonferroni correction for multiple pairwise testing. Many functional markers were found to be significantly more abundant in specific sites; more will be reported in each of the following sections (Figure 1, Table S3). These contrasting abundances of functional marker genes suggest significant differences in the metabolisms of microbial communities across lake environments.

**Figure 1.**
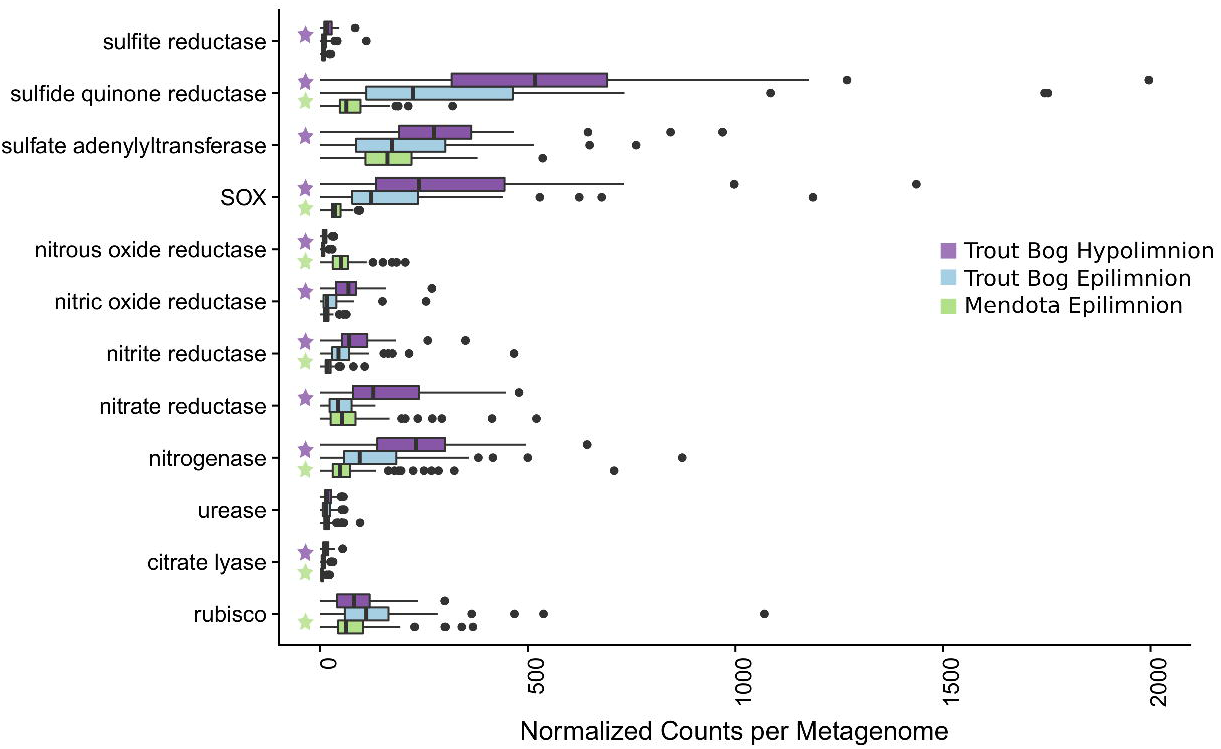
Analysis of marker gene abundances reveals differences between lakes and layers. To assess potential differences in microbial metabolisms in our study sites, we predicted open reading frames in unassembled metagenomes using Prodigal and compared the resulting ORFs to a custom database of metabolic marker genes using BLAST. In these boxplots, significant differences in numbers of gene hits between sites was tested using a pairwise Wilcoxon rank sum test with a Bonferroni correction; significance was considered to be p < 0.05. 94 metagenomes were tested for Lake Mendota, while 47 metagenomes were tested in each layer of Trout Bog. Significant differences between the Trout Bog and Lake Mendota epilimnia and between the Trout Bog epilimnion and hypolimnion are indicated by a green or a purple star, respectively. Significant differences between the Trout Bog hypolimnion and the Lake Mendota epilimnion were not tested, as the large number of variables differing in these sites makes the comparison less informative. This analysis revealed differences in the number of marker genes observed by lake for many metabolic processes involved in carbon, nitrogen, and sulfur cycling. LEfSe results for each gene are available in Data S3, and p-values of markers described in Figure 1 and elsewhere in the text are reported in Table S3.

### How Representative are the MAGs?

To identify the phylogenies of the microbes carrying marker genes and the co-occurrences of marker genes within the same population genomes, we used metagenome-assembled genomes (MAGs) from each metagenomic time series to predict metabolic pathways based on genomic content. A total of 193 medium to high quality bacterial MAGs were recovered from the three combined time series metagenomes in Trout Bog and Lake Mendota: 99 from Lake Mendota, 31 from Trout Bog’s epilimnion, and 63 from Trout Bog’s hypolimnion (Data S4). These population genomes ranged in estimated completeness from 50 to 99% based on CheckM estimates (Parks et al., 2015). Several MAGs from Trout Bog’s epilimnion and hypolimnion appeared to belong to the same population based on average nucleotide identities greater than 99% calculated using DOE JGI’s ANI calculator (Data S6) (Varghese et al., 2015). This is likely because assembly and binning were carried out separately for each thermal layer, even though some populations were present throughout the water column. To assess the diversity of our MAGs, we constructed an approximate maximum likelihood tree of all the MAGs in FastTree (Price, Dehal & Arkin, 2010) using whole genome alignments (Figure S1). The tree is not intended to infer detailed evolutionary history, but to provide an overall picture of similarity between genomes. MAGs recovered are a diverse set of genomes assigned to taxa typically observed in freshwater.

The phylum-level assignments of our MAGs largely matched the classifications of 16S rRNA gene amplicon sequencing results averaged across the time series, consistent with a higher likelihood of recovering MAGs from the most abundant populations in the community (Figure S2, Data S5). However, some taxa, including *Tenericutes, Ignavibacteria, Epsilonproteobacteria*, and *Chlamydiae*, were represented by MAGs but not identified in the 16S gene amplicon datasets. *Chlorobi* was overrepresented by MAG coverage compared to 16S rRNA gene counts, while *Proteobacteria* was overrepresented by 16S rRNA gene counts compared to MAG coverage. These discrepancies could be explained by bias in the 16S primer sets (Hong et al., 2009) difference in *rRNA* copy number, or assembly bias in MAG recovery. The observed taxonomic compositions are consistent with other 16S-based studies from these lakes (Hall et al., 2017; Linz et al., 2017). The detection of similar phyla using both methods suggests that our MAGs are representative of the resident microbial communities.

### Nitrogen Cycling

Nitrogen availability is an important factor structuring freshwater microbial communities. To see if there were differences in nitrogen cycling between different lake environments, we analyzed nitrogen-related marker genes and the MAGs containing nitrogen cycling pathways. We discovered significant differences in the abundances of marker genes, along with phylogenetic differences in the populations containing these pathways.

To identify differences in nitrogen fixation between sites, we analyzed marker genes encoding nitrogenase subunits. Genes encoding for nitrogenase were observed most frequently in metagenomes from Trout Bog’s hypolimnion, followed by the Trout Bog’s epilimnion, and lastly by Lake Mendota’s epilimnion (Figure 1, Table S3). The nitrogenase enzyme is inhibited by oxygen, which could explain the higher abundance of nitrogenase in Trout Bog’s anoxic hypolimnion. We further analyzed MAGs predicted to fix nitrogen and found differences in the taxonomy of putative diazotrophs between the two ecosystems (Figure 2, Figure S1). In Lake Mendota, two thirds of MAGs encoding the nitrogen fixation pathway were classified as *Cyanobacteria*, while the other third was assigned to *Betaproteobacteria* and *Gammaproteobacteria*. Although not all *Cyanobacteria* fix nitrogen, previous measurements of nitrogen fixation in Lake Mendota found a strong correlation between this pathway and the *Cyanobacteria Aphanizomenon* (Beversdorf, Miller & McMahon, 2013). MAGs containing genes encoding nitrogen fixation were more phylogenetically diverse in Trout Bog and included *Deltaproteobacteria, Gammaproteobacteria, Epsilonproteobacteria, Acidobacteria, Verrucomicrobia, Chlorobi*, and *Bacteroidetes*. The increased diversity of diazotrophs in Trout Bog compared to Lake Mendota suggests that nitrogen fixation genes may be horizontally transferred with populations in Trout Bog.

**Figure 2.**
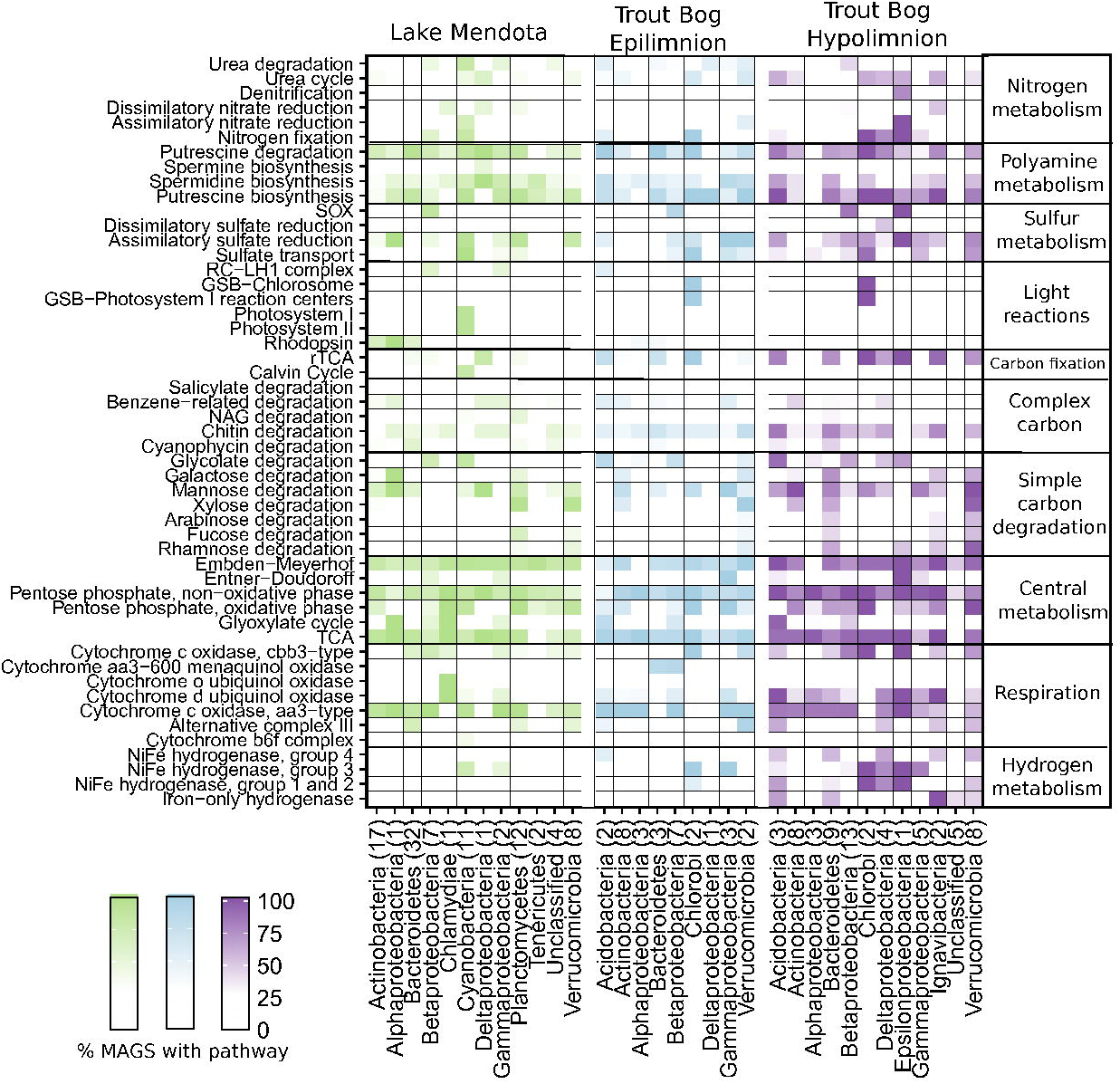
Metabolisms in Lake Mendota and Trout Bog. Metabolic pathways were predicted for all MAGs based on their gene content. At least 50% of enzymes in a pathway must have been encoded in the genome for a pathway to be considered present, as well as encoding enzymes unique to or required for a pathway. Putative pathway presence was aggregated by lake and phylum. This analysis can link potential functions identified in the metagenomes to taxonomic groups that may perform those functions. For example, MAGs with putative pathways for carbon fixation also likely fix nitrogen in both lakes. Similar, putative degradation pathways for rhamnose, fucose, and galactose were frequently encoded in the same MAGs. *Proteobacteria* was split into classes due to the high diversity of this phylum.

To identify differences in denitrification, we analyzed marker genes for denitrification, including reductases for nitrous oxide, nitric oxide, nitrite, and nitrate. These denitrification genes had a similar trend as the nitrogen fixation genes; they were observed most frequently in metagenomes from the Trout Bog hypolimnion, with the exception of nitrous oxide reductase, which was most frequently found in Lake Mendota. This trend could stem from denitrification also requiring a reductive, low oxygen environment. Urease, another nitrogen cycling marker gene, was not found significantly more often in any site. We further analyzed putative denitrification pathways in our MAGs and found that they were observed at similar frequencies in population genomes from all environments (Figure 2). Urea degradation pathways were also predicted in MAGs from both lakes, which is consistent with research showing that urea is a common nitrogen source for bacteria in multiple freshwater environments (Remsen, Carpenter & Schroeder, 1972; Jorgenson et al., 1998; Berman & Bronk, 2003).

To explore the importance of polyamines in the freshwater nitrogen cycle, we analyzed genes encoding the biosynthesis and degradation of polyamines such as spermidine and putrescine. We predicted that 94% of MAGs could synthesize polyamines, and 87% could degrade polyamines. These genes were prevalent in many diverse MAGs from both lakes, including *Actinobacteria* as has been previously observed (Ghylin et al., 2014; Hamilton et al., 2017). While there is some evidence for the importance of polyamines in aquatic systems (Mou et al., 2011), the ecological role of these compounds in freshwater is not fully resolved.

Polyamines are known to play a critical but poorly understood role in bacterial metabolism (Igarashi & Kashiwagi, 1999), and the exchange of these nitrogen compounds between populations may be a factor structuring freshwater microbial communities. Polyamines can also result from the decomposition of amino acids, so higher trophic levels such as fish or zooplankton may provide an additional source (Al Bulushi et al., 2009). The frequent appearance of polyamine-related pathways in our MAGs lends support to the hypothesis that these compounds are important parts of the dissolved organic nitrogen and carbon pool in freshwater.

To identify signatures of nitrogen limitation at the genomic level, we analyzed biases in amino acid use in our MAGs (Data S4) (Acquisti, Kumar & Elser, 2009; Bragg & Wagner, 2009). For this analysis, genomes from the Trout Bog layers were considered together due to the previously mentioned overlap in recovered genomes. We observed that on average, MAGs from Trout Bog encoded amino acids with 1% less nitrogen than MAGs from Lake Mendota. Although this difference is small, it was significant using a Wilcoxon rank sum test (p = 0.02). The observed amino acid bias suggests that conditions in Trout Bog may lead to stronger selection for nitrogen poor proteins than in Lake Mendota. Differences in the compositions of the nitrogen pools in these lakes may also contribute to the observed differences in the distributions of nitrogen cycling marker genes. Lake Mendota receives large amounts of nitrate runoff from the surrounding agricultural landscape, while Trout Bog receives nitrogen in more complex forms (e.g. *Sphagnum-derived* organic nitrogen), and the microbial community competes for nitrogen with the surrounding plant community.

### Sulfur Cycling

Sulfur is another essential element in freshwater that is cycled between oxidized and reduced forms by microbes. Our marker gene analysis demonstrated that genes encoding for sulfide:quinone reductase (for sulfide oxidation) and the sox pathway (for thiosulfate oxidation) were significantly more abundant in Trout Bog compared to Lake Mendota, with no significant differences between the layers of Trout Bog (Figure 1, Table S3). Genes encoding for sulfite reductases were the least abundant sulfur cycling marker genes in all sites. Dissimilatory sulfite reductase was observed only in MAGs from Trout Bog, especially those classified as *Chlorobiales*. Because this enzyme is thought to operate in reverse in green sulfur-oxidizing phototrophs such as *Chlorobiales* (Holkenbrink et al., 2011), this may indicate an oxidation process rather than a reductive sulfur pathway. Assimilatory sulfate reduction was the most common sulfur-related pathway identified in the MAGs (Figure 2).

We observed assimilatory sulfate reduction more frequently than dissimilatory sulfate reduction, suggesting that in these populations, sulfate is more commonly used for biosynthesis, while reduced forms of sulfur are used as electron donors for energy mobilization. This is in contrast to marine systems, where sulfate reduction holds a central role as an energy source for organotrophic energy acquisition (Bowles et al., 2014), although sulfate reduction could also be occurring in Lake Mendota’s hypolimnion. Sulfur oxidation pathways were observed in MAGs classified as *Betaproteobacteria* from both lakes and *Epsilonproteobacteria* in Trout Bog’s hypolimnion.

### Phototrophy

Primary production (the coupling of photosynthesis and carbon fixation) is a critical component of the freshwater carbon cycle. To identify differences in routes of primary production between freshwater environments, we compared marker genes for carbon fixation across sites. RuBisCO (ribulose-1,5-bisphosphate carboxylase/oxygenase), the marker gene for carbon fixation via the Calvin-Benson-Bassham (CBB) pathway, was most frequently observed in Trout Bog’s epilimnion (Figure 1, Table S3). In contrast, citrate lyase, the marker gene for the reverse TCA cycle, was observed most frequently in Trout Bog’s hypolimnion.

We next assessed the MAGs for photoautotrophy, expecting to find differences between our two study sites based on the observed contrasts in the functional marker gene analysis (Figure 2). In Lake Mendota, the majority of MAGs encoding phototrophic pathways were classified as *Cyanobacteria*. These populations contained genes encoding enzymes in the CBB pathway. In Trout Bog, most MAGs encoding phototrophy were classified as *Chlorobium clathratiforme*, a species of *Chlorobiales* widespread in humic lakes (Karhunen et al., 2013). The *Chlorobiales* MAGs in Trout Bog contained genes encoding citrate lyase and other key enzymes in the reductive tricarboxylic acid (TCA) cycle, an alternative carbon fixation method commonly found in green sulfur bacteria such as *Chlorobi* (Kanao et al., 2002; Tang & Blankenship, 2010). As *Chlorobium* is a strictly anaerobic lineage, the presence of citrate lyase in these populations may explain why this gene was observed more frequently in metagenomes from Trout Bog’s hypolimnion. These photoautotrophs from both lakes also contained genes potentially encoding nitrogen fixation. The co-occurrence of fixation pathways in these populations are especially interesting given their relatively high abundance in their respective lakes.

The reductive TCA cycle is the only carbon fixation pathway known to be active in cultured representatives of *Chlorobiales*, but we found genes annotated as the RuBisCO large subunit (rbcL) were observed in some of the *Chlorobiales* MAGs. Homologs of *rbcL* have been previously identified in isolates of *Chlorobium*, and were associated with sulfur metabolism and oxidative stress (Hanson & Tabita, 2001). Inspection of the neighborhoods of genes annotated as *rbcL* in the *Chlorobiales* MAGs revealed genes putatively related to rhamnose utilization, LPS assembly, and alcohol dehydrogenation, but no other CBB pathway enzymes. Given this information, it seems likely that this *rbcL* homolog encodes a function other than carbon fixation in the *Chlorobiales* MAGs.

The potential for photoheterotrophy via the aerobic anoxygenic phototrophic pathway was identified in several MAGs from all lake environments, especially from epilimnia, based on the presence of genes annotated as *pufABCLMX, puhA*, and *pucAB* encoding the core reaction center RC-LH1 (Martinez-Garcia et al., 2012). *Betaproteobacteria* and *Gammaproteobacteria*, particularly MAGs classified as *Burkholderiales*, most often contained these genes, although they were not broadly shared across the phylum (Figure 2). As aerobic anoxygenic phototrophy has previously been associated with freshwater *Proteobacteria* (Martinez-Garcia et al., 2012), these results are not surprising. Unexpectedly, an *Acidobacteria* MAG from the Trout Bog epilimnion also contained genes suggesting aerobic anoxygenic phototrophy.

Another form of photoheterotrophy previously identified in freshwater is the use of light-activated proteins such as rhodopsins (Martinez-Garcia et al., 2012). We observed genes encoding rhodopsins in MAGs from each lake environment, but more frequently in *Actinobacteria* and *Bacteroidetes* MAGs from Lake Mendota (Figure 2). Trout Bog, especially the hypolimnion, harbored fewer, less diverse MAGs encoding rhodopsins than those from Lake Mendota.

### Complex Carbon Degradation

Biopolymers in freshwater can be either autochthonous (produced within the lake, ex. algal polysaccharides) or allochthonous (imported from the surrounding landscape, ex. cellulose). Organic carbon in freshwater is often classified as either autochthonous or allochthonous carbon, but this distinction has little relevance for organotrophic bacteria. For example, there is substantial overlap in the molecular composition of algal exudates, cellulose degradation intermediates, and photochemical degradation products (Bertilsson & Tranvik, 1998; Ramanan et al., 2015). One-carbon compounds such as methane are produced in the lake (therefore autochthonous), but they are also produced from decomposition of allochthonous carbon. We therefore found it more informative to categorize the carbon degradation pathways observed in our dataset by type of metabolism rather than carbon origin.

Degradation of high-complexity, recalcitrant carbon compounds requires specialized enzymes, but a wide availability of these compounds can make complex carbon degradation an advantageous trait. One way to predict the ability to degrade high-complexity carbon in microbial populations is by identifying genes annotated as glycoside hydrolases (GHs), which encode enzymes that break the glycosidic bonds found in complex carbohydrates. A previous study of *Verrucomicrobia* MAGs from our dataset found that the profiles of GHs differed between Lake Mendota and Trout Bog, potentially reflecting the differences in available carbon sources (He et al., 2017). Here, we expanded this analysis of glycoside hydrolases to all of the MAGs in our dataset to identify differences in how populations from our two study sites degrade complex carbohydrates.

We calculated the coding density of GHs, defined as the percentage of coding regions in a MAG annotated as a GH to identify differences in carbon metabolism between MAGs from different lake environments (Figure 3). Our GH coding density metric was significantly correlated with the diversity of GHs identified (r^2^= 0.39, p = 4.5×10^−8^), which is an indicator of the number of substrates an organism can utilize. The MAGs with the highest GH coding densities were classified as *Bacteroidales, Ignavibacteriales, Sphingobacteriales*, and *Verrucomicrobiales* from Trout Bog’s hypolimnion. Two of these orders, *Sphingobacteriales* and *Verrucomicrobiales*, also contained MAGs with high GH coding densities in Lake Mendota and Trout Bog’s epilimnion. There were several additional orders with high GH coding density that were unique to Lake Mendota, including *Mycoplasmatales (Tenericutes), Cytophagales (Bacteroidetes), Planctomycetales (Planctomycetes)*, and *Puniceicoccales (Verrucomicrobia)*. In concordance with their ability to hydrolytically degrade biopolymers to sugars, MAGs with high GH coding densities also contained putative degradation pathways for a variety of sugars (Figure 2).

**Figure 3.**
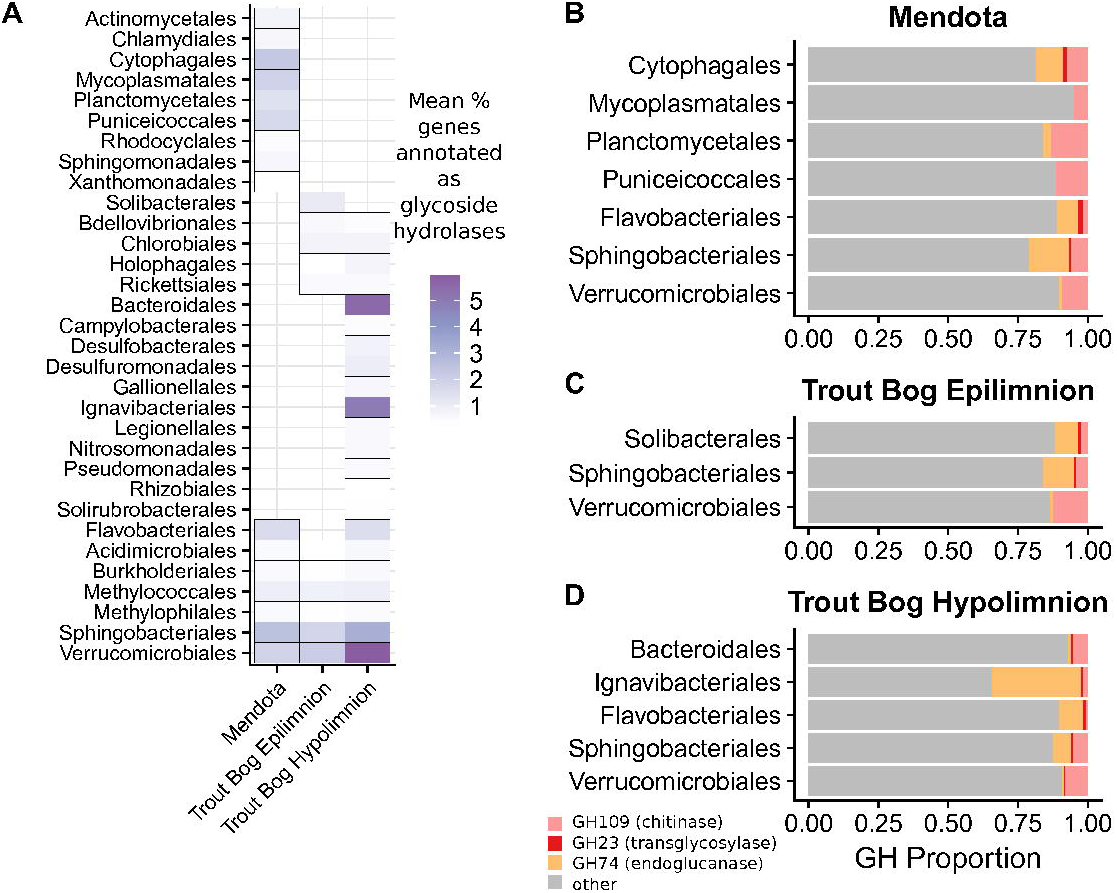
Glycoside hydrolase content in the MAGs. Annotations of GHs were used as an indication of complex carbon degradation. Genes potentially encoding GHs were identified and assigned CAZyme annotations using dbCAN. GH coding density was calculated for each MAG and averaged by order and lake (A). While a few orders contained genes encoding glycoside hydrolases in all three sites, many orders were unique to each site. The orders with the highest coding density were all found in the Trout Bog hypolimnion. Glycoside hydrolase diversity, an indicator of the range of substrates an organism can degrade, was significantly correlated with coding density (r2 = 0.38, p = 4.5×10-8). Within MAGs with high glycoside hydrolase density, three families appeared most frequently - GH74, GH109, and GH23, although these abundances may be method-dependent (He et al., 2017) (B-D). *Proteobacteria* was split into classes due to the high diversity of this phylum.

We identified genes encoding for several GH families in MAGs from all lake environments. Starting with the most frequently observed in MAGS from all sites, these included GH109 (alpha-N-acetylgalactosaminidase), GH74 (endoglucanase), and GH23 (soluble lytic transglycosylase). However, previous research has found that the abundance of genes annotated as GH109 by dbCAN may be an overestimate of this gene family (He et al., 2017); therefore, we prefer not to speculate on the relative importance of GH family annotations in our MAGs based on observation frequency. Lake Mendota contained unique GHs belonging to the family GH13 (alpha-glucoside). The only unique GH found in Trout Bog’s epilimnion was GH62, a putative arabinofuranosidase. Trout Bog’s hypolimnion contained many more unique enzymes, the most abundant of which were GH129 (alpha-N-acetylgalactosaminidase), GH89 (alpha-N-acetylglucosaminidase), GH43_12 (xylosidase/arabinosidase), GH44 (beta-mannanase/endo-beta-1,4-glucanase), GH66 (dextranase), and GH67 (alpha-glucuronidase). The increased diversity of these genes found in Trout Bog’s hypolimnion suggests differences between the GH profiles, which could be correlated to differing diversity and complexity of the available organic carbon.

### Central Metabolism and Simple Carbon Degradation

Freshwater microbes are exposed to a great variety of low-complexity carbon sources such as carbohydrates, carboxylic acids, and one-carbon (C1) compounds. The central metabolic pathways shared by most living cells are often an entry point for the least complex carbon compounds. The specific routing of central metabolism may therefore reveal how low complexity carbon compounds are used. Genes encoding enzymes in the glyoxylate cycle, a truncated version of the TCA cycle that is used to produce biosynthetic intermediates and bypass decarboxylation steps, were observed in *Alphaproteobacteria* and *Chlamydiae* in Lake Mendota and *Acidobacteria* and *Betaproteobacteria* in Trout Bog. This may indicate an adaptation to reduce carbon demand in these populations.

Oxidative phosphorylation is an important part of central metabolism for aerobic bacteria, so we investigated the types of cytochrome oxidases encoded in our MAGs (Figure 2). Cytochrome c oxidases, both aa3- and cbb3-type, were widespread in all three lake environments and frequently co-occurred within MAGs. aa3-type cytochromes are associated with high oxygen concentrations and cbb3-type cytochromes are associated with low oxygen concentrations (Gong et al., 2018), so the presence of genes encoding both types suggests the flexibility to operate under a range of oxygen concentrations. Of the quinol-based cytochrome oxidases, genes encoding cytochrome d oxidase were most often observed in MAGs from Trout Bog’s hypolimnion, while cytochrome aa3-600 was found only in MAGs classified as *Bacteroidetes* and *Betaproteobacteria* from Trout Bog’s epilimnion. Cytochrome o oxidase was observed only in a *Chlamydia* MAG from Lake Mendota. Alternative complex III was identified in MAGs of *Verrucomicrobia* in all sites, in *Acidobacteria* from Trout Bog (both layers), and in *Bacteroidetes* and *Planctomycetes* from Lake Mendota.

Similarly, hydrogen metabolism can influence other aspects of a microbe’s nutrient usage. Iron-only hydrogenases were found primarily in MAGs from Trout Bog’s hypolimnion (Figure 2, Table S3), consistent with their previously identified presence in anaerobic, often fermentative bacteria (Peters et al., 2015) and the higher observations of marker genes for iron-only hydrogenases in the hypolimnion site. Genes encoding [Ni-Fe] hydrogenases of groups 1 and 2, involved in hydrogen uptake, sensing, and nitrogen fixation, were found at significantly different frequency in all sites with the exceptions of group 2a in Lake Mendota and Trout Bog’s epilimnion and group 2b in both layers of Trout Bog. Genes encoding these hydrogenases were widespread in MAGs from Trout Bog’s hypolimion, found only in *Chlorobiales* MAGs in Trout Bog’s epilimnion, and rarely observed in MAGs from Lake Mendota. Group 3 [Ni-Fe] hydrogenases were detected differentially at each site dependent on their subtype and were identified in MAGs belonging to *Cyanobacteria* and *Chlorobiales* in both lakes. This finding is consistent with the proposed function of Group 3d, which is to remove excess electrons produced by photosynthesis. Group 4 [Ni-Fe] hydrogenases were not observed significantly more or less in any site.

Low molecular weight carbohydrates such as glucose, fucose, rhamnose, arabinose, galactose, mannose, and xylose may be derived either from algae or from cellulose degradation (Giroldo, Augusto & Vieira, 2005; Ramanan et al., 2015). To understand how these compounds are used by freshwater populations, we analyzed putative sugar degradation pathways in our MAGs. Genes encoding the pathway for mannose degradation, which feeds into glycolysis, appeared frequently in both lakes. Genes encoding the degradation of rhamnose and fucose, whose pathways converge to enter glycolysis and produce pyruvate, were frequently found within the same MAGs (including members of *Planctomycetes* and *Verrucomicrobia* from Lake Mendota, and members of *Bacteroidetes, Ignavibacteria*, and *Verrucomicrobia* from Trout Bog). Putative pathways for galactose degradation were often observed in these same MAGs. Xylose is a freshwater sugar which has already been identified as potential carbon source for streamlined Actinobacteria (Ghylin et al., 2014); we confirmed this in our MAGs, and found that *Bacteroidetes, Planctomycetes*, and *Verrucomicrobia* from Lake Mendota and *Bacteroidetes* and *Verrucomicrobia* from Trout Bog were additional potential xylose degraders. Genes for the degradation of glycolate, an acid produced by algae and consumed by heterotrophic bacteria (Paver & Kent, 2017), were identified in *Cyanobacteria* and *Betaproteobacteria* MAGs from Lake Mendota and in *Acidobacteria, Verrucomicrobia, Alpha-, Beta-, Gamma-*, and *Epsilonproteobacteria* MAGs from Trout Bog.

Methylotrophy, the ability to grow solely on C1 compounds such as methane or methanol, was predicted in MAGs from both Trout Bog and Lake Mendota. Putative pathways for methanol degradation were found in MAGs classified as *Methylophilales* (now merged with *Nitrosomonadales* (Boden, Hutt & Rae, 2017)) and *Methylotenera*, while *Methylococcales* MAGs were potential methane degraders based on the presence of genes encoding methane monooxygenase. *Methylococcales* MAGs from Trout Bog also encoded the pathway for nitrogen fixation, consistent with reports of nitrogen fixation in cultured isolates of this taxon (Bowman, Sly & Stackebrandt, 1995). The *Methylophilales* MAGs also likely degrade methylamines, based on the presence of genes encoding the N-methylglutamate pathway or the tetrahydrofolate pathway (Latypova et al., 2010). Methylotrophy in cultured freshwater isolates from these taxa is well-documented (Kalyuzhnaya et al., 2011; Salcher et al., 2015); however, genes encoding methanol degradation were also identified in MAGs classified as *Burkholderiales* and *Rhizobiales* from Trout Bog. Given the rapid rate at which we are discovering methylotrophy in microorganisms not thought to be capable of this process, identifying potential new methylotrophs in freshwater is intriguing, but not surprising (Chistoserdova, Kalyuzhnaya & Lidstrom, 2009).

### Using MAGs to track population abundances over time

Our metagenomes comprise a time series, so we can use MAG coverage and the number of marker gene hits as proxies for abundance over time. As an example, we analyzed abundance data for *Cyanobacteria*, known to be highly variable over time in Lake Mendota (Figure 4, A-E). We found that one *Cyanobacteria* MAG in each year was substantially more abundant than the rest; this single MAG only is plotted for each year. Since our analysis of the diversity of MAGs containing nitrogenases showed a strong association between nitrogen fixation and *Cyanobacteria* in Lake Mendota, we hypothesized that the number of hits to the most abundant marker genes encoding nitrogenase subunits over time would be correlated to the abundance of the most abundant *Cyanobacteria* MAG in each year (Figure 4, F-J). This hypothesis was partially supported. Two of the marker genes, TIGR1282 (*nifD*) and TIGR1286 (*nifK* specific for molybdenum-iron nitrogenase), correlated with the *Cyanobacteria* MAG abundance more frequently than the third, TIGR1287 (*nifH*, common among different types of nitrogenases). Significant correlations (p < 0.05) were only detected in 2008, 2011, and 2012. The strength of these correlations suggests that in three out of the five years in our Lake Mendota time series, a single *Cyanobacteria* population produced most genes encoding nitrogenase subunits. In the other two years, it is possible that other diazotrophic populations were more abundant, or that the nitrogenase subunits were derived from populations that did not assemble into MAGs. These two years were also unusual in our time series - in 2008, extreme flooding events led to large *Cyanobacteria* blooms (Beversdorf et al., 2015) and in 2009, the invasive spiny water flea population drastically increased in Lake Mendota (Walsh, Munoz & Vander Zanden, 2016). Still, our time series analysis demonstrates the utility of our datasets in linking metabolic function to specific taxonomic groups.

**Figure 4.**
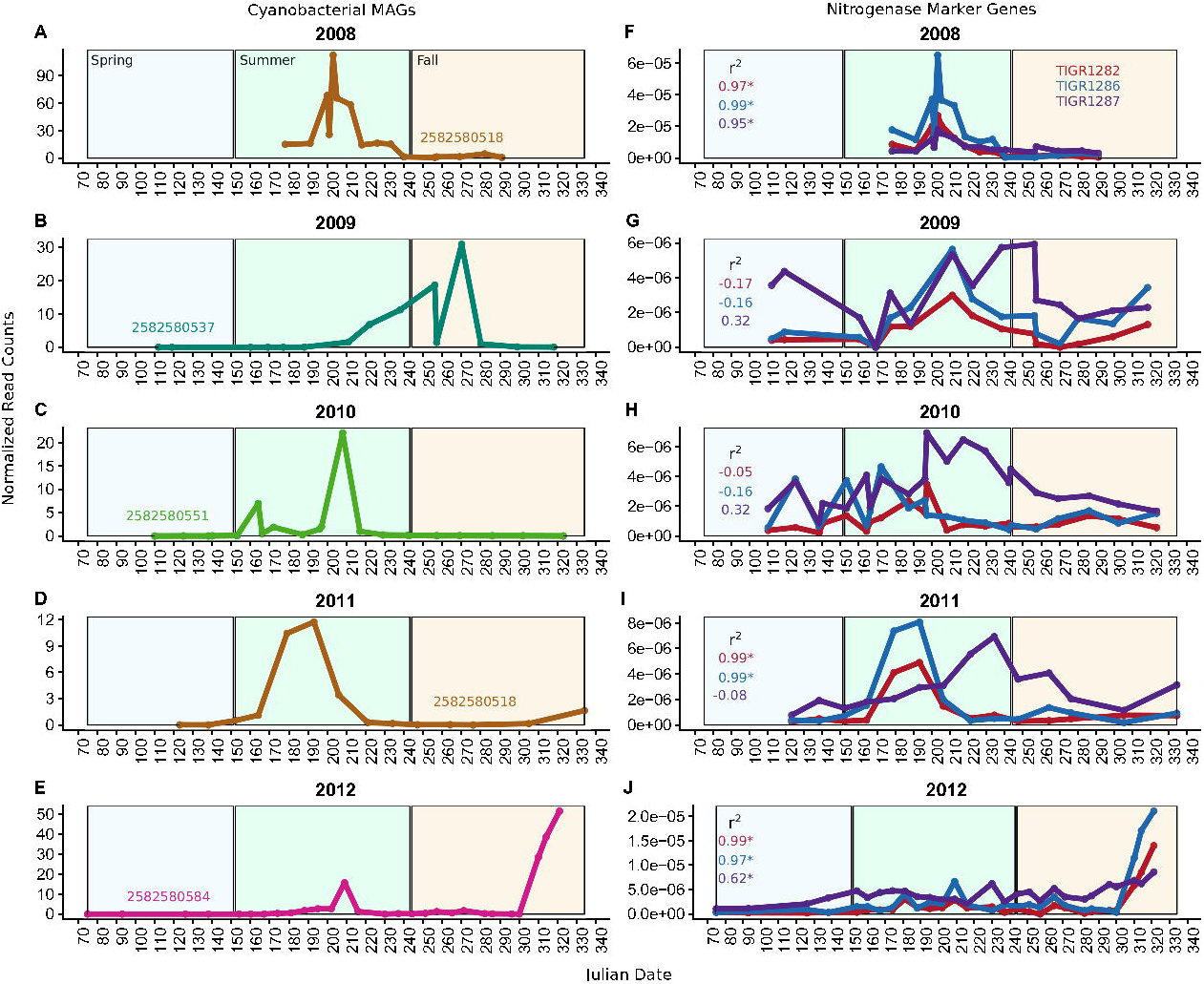
Cyanobacteria and nitrogen fixation over time. To approximate the abundance of populations over time, we mapped metagenomic reads back to the MAGs. The number of BLAST hits of marker genes in the metagenomes was used as a proxy for gene abundance. Counts were normalized by metagenome size, and in the case of the MAGs, genome length. Data from *Cyanobacteria* MAGs and nitrogen fixation marker genes are shown here. Colored numbers on panels A, C, E, G, and I indicate the IMG OID of the most abundant MAG in that year of data, plotted here. The marker genes used were TIGR1282, TIGR1286, and TIGR1287, encoding subunits of Mo-Fe nitrogenase; these were the most frequently observed nitrogenase markers in the Lake Mendota metagenomes. Significantly correlated trends over time were observed in the MAGs and the nitrogenase marker genes in 2008, 2011, and 2012. This suggests that nitrogen fixation is driven by these particular MAGs in those years, and is consistent with our result indicating that genes encoding nitrogen fixation were found in these MAGs. The lack of significant correlations in other years may be due to contributions from unassembled populations or more even abundances of other diazotrophic populations in that year.

## Conclusions

Our analysis of functional marker genes indicated significant differences in microbial nutrient cycling between Lake Mendota’s epilimnion, Trout Bog’s epilimnion, and Trout Bog’s hypolimnion. By combining these results with metabolic pathway prediction in MAGs, we identified taxa encoding these metabolisms and co-occurrence of pathways within MAGs. We found that phototrophy, carbon fixation, and nitrogen fixation co-occurred within the abundant phototrophs *Cyanobacteria* in Lake Mendota and *Chlorobiales* in Trout Bog. In Lake Mendota, nitrogen fixation was predominantly associated with *Cyanobacteria*, but it was not associated with any particular taxon in Trout Bog. In the sulfur cycle, we observed assimilatory pathways more frequently than dissimilatory pathways in the MAGs, suggesting a bias towards using sulfur compounds for biosynthesis rather than as electron donors. We found the greatest density and diversity of genes annotated as GHs in the Trout Bog hypolimnion, potentially indicating a greater reliance on complex carbon sources in this environment. Our combination of functional marker gene analysis and MAG pathway prediction provided insight into the complex metabolisms underpinning freshwater communities and how microbial processes scale to ecosystem functions.

## Acknowledgments

We thank the North Temperate Lakes and Lake Mendota Microbial Observatory field crews, UW-Trout Lake Station, the UW Center for Limnology, and the Global Lakes Ecological

Observatory Network for field and logistical support. We acknowledge efforts by many McMahon laboratory undergraduate students and technicians whose work has been related to sample collection and DNA extraction. We thank Emily Stanley and Joshua Hamilton for insightful comments on an early draft of this manuscript. Finally, we personally thank the individual program directors and leadership at the National Science Foundation for their commitment to continued support of long-term ecological research.

We thank the Joint Genome Institute for supporting this work through the Community Sequencing Program (CSP 394), performing the bioinformatics, and providing technical support. The work conducted by the U.S. Department of Energy Joint Genome Institute, a DOE Office of Science User Facility, is supported by the Office of Science of the U.S. Department of Energy under Contract No. DE-AC02-05CH11231. K.D.M. acknowledges funding from the United States National Science Foundation Microbial Observatories program (MCB-0702395), the Long Term Ecological Research Program (NTL-LTER DEB- 1440297), and an INSPIRE award (DEB-1344254). A.M.L. was supported by a pre-doctoral fellowship provided by the University of Wisconsin – Madison Department of Bacteriology and by the National Science Foundation Graduate Research Fellowship Program under grant no. DGE-1256259 during this research. This material is also based upon work that supported by the National Institute of Food and Agriculture, U.S. Department of Agriculture (Hatch Project 1002996).

## Figure and Table Legends

**Table 1. Characteristics of Lake Mendota and Trout Bog.** Water from Lake Mendota and Trout Bog was sampled weekly during the ice-free periods using an integrated water column sampler and filtered for DNA using a 0.22 micron filter. Metagenomic sequencing was performed on DNA extracted from filters collected in 2008-2012 from Lake Mendota and in 2007-2009 from Trout Bog. The epilimnion (upper thermal layer) was sampled in both lakes, while the hypolimnion (bottom thermal layer) was sampled only in Trout Bog. Chemistry data were collected by NTL-LTER from depth discrete samples taken from 0 and 4 m for Lake Mendota, 0 m for the Trout Bog Epilimnion, and 3 and 7 m for the Trout Bog Hypolimnion. Values reported here are the means of all measurements in the sampling time span for each lake, with standard deviations reported in parentheses.

## Supplemental Legends

**Table S1. Additional chemical measurements in our study sites.** Additional chemistry data were collected by NTL-LTER (http://lter.limnology.wisc.edu) from depth discrete samples taken from 0 and 4 m for Lake Mendota, 0 m for the Trout Bog Epilimnion, and 3 and 7 m for the Trout Bog Hypolimnion. Values reported here are the means of all measurements in the sampling time span for each lake, with standard deviations reported in parentheses.

**Data S1. IMG Genome ID numbers and information about metagenomes used in this study.** This dataset includes information about the metagenomes used in this study including date collected, size in reads and base pairs, and their IMG Genome IDs (IMG Taxon ID).

**Data S2. Functional marker genes used in this study.** This dataset lists the TIGRFAM, COG, or PFAM IDs of sequences used as functional marker genes to analyze how gene content differs by site.

**Table S2. Statistics from genome assembly and binning.** Metagenomic samples were pooled by lake and layer to allow time-resolved binning. The time series in Lake Mendota spans 2008-2012, while the Trout Bog time series spans 2007-2009. The large amount of DNA assembled produced just under 200 medium to high quality metagenome-assembled genomes.

**Data S3. Results of LEfSe analysis on functional marker genes**. The program LEfSe was used to detect significant differences in gene content between our study sites. The distinguish feature of LEfSe, the LDA effect score, is listed for each marker gene in this dataset.

**Table S3. P-values of marker gene distributions between sites.** A Wilcoxon rank sum test was used to non-parametrically test for significant differences in functional marker gene distributions between our study sites. P-values of less than 0.05 are considered significant.

**Data S4. MAG metadata.** Information about the completeness, size, and taxonomy of our MAGs, as well as their IMG OIDs, are presented here. Amino acid use was calculated based on the average number of nitrogen atoms translated gene sequences.

**Data S5. 16S rRNA amplicon sequencing of our samples.** 16S sequencing was performed over the time series to assess community composition in our study sites. The resulting OTU tables and taxonomic classifications are presented here.

**Figure S1. Tree of diversity and nitrogen fixation in our MAGs**. To visualize the diversity of our MAGs, phylogenetic marker genes were extracted from each MAG and aligned using Phylosift. An approximate maximum-likelihood tree based on these alignments was constructed using FastTree. The potential for nitrogen fixation based on gene content is indicated on the branch tips.

**Figure S2. How representative are the MAGs of the microbial communities?** The community composition observed via 16S rRNA gene amplicon sequencing (A) and inferred using the proportions of reads from the same metagenomic time series samples that mapped to set of MAGs affiliated with major phyla (B). MAGs were classified using Phylosift, while 16S sequences were classified to the phylum level. Numbers above bars indicating abundances greater than the limit of the y-axis. The 16S V6-V8 region was targeted in Trout Bog, while the V4 region was targeted in Lake Mendota. *Proteobacteria* was split into classes due to the high diversity of this phylum. Although proportions vary, similar taxonomic groups are observed using both approaches. Differences are likely due to a combination of primer and assembly biases. However, similar phyla were detected using both methods, suggesting that our MAG datasets are representative of their communities.

**Data S6. Average nucleotide identity between MAGs.** Average nucleotide identity (ANI) was calculated between all MAGs in our dataset. MAGs with extremely high ANIs (>97%) are likely from the same populations. An ANI value of “0” indicates that no portions of the genomes aligned.

## References

Acquisti C., Kumar S., Elser JJ. 2009. Signatures of nitrogen limitation in the elemental composition of the proteins involved in the metabolic apparatus. Proceedings of the Royal Society B: Biological Sciences 276:2605–2610. DOI: 10.1098/rspb.2008.1960.

Allgaier M., Grossart H-P. 2006. Diversity and Seasonal Dynamics of Actinobacteria Populations in Four Lakes in Northeastern Germany. Applied and Environmental Microbiology 72:3489–3497. DOI: 10.1128/AEM.72.5.3489.

Anantharaman K., Brown CT., Hug LA., Sharon I., Castelle CJ., Probst AJ., Thomas BC., Singh A., Wilkins MJ., Karaoz U., Brodie EL., Williams KH., Hubbard SS., Banfield JF. 2016. Thousands of microbial genomes shed light on interconnected biogeochemical processes in an aquifer system. Nature Communications 7:1–11. DOI: 10.1038/ncomms13219.

Azam F., Fenchel T., Field JG., Gray JC., Meyer-Reil LA., Thingstad F. 1983. The ecological role of water-column microbes in the sea. Marine Ecology Progress Series 10:257–264. DOI: 10.3354/meps010257.

Bendall ML., Stevens SLR., Chan L., Malfatti S., Schwientek P., Tremblay J., Schackwitz W., Martin J., Pati A., Bushnell B., Froula J., Kang D., Tringe SG., Bertilsson S., Moran MA., Shade A., Newton RJ., McMahon KD., Malmstrom RR. 2016. Genome-wide selective sweeps and gene-specific sweeps in natural bacterial populations. The ISME Journal 10:1589–1601. DOI: 10.1038/ismej.2015.241.

Berman T., Bronk DA. 2003. Dissolved organic nitrogen: A dynamic participant in aquatic ecosystems. Aquatic Microbial Ecology 31:279–305. DOI: 10.3354/ame031279.

Bertilsson S., Tranvik LJ. 1998. Photochemically produced carboxylic acids as substrates for freshwater bacterioplankton. Limnology and Oceanography 43:885–895. DOI: 10.4319/lo.1998.43.5.0885.

Beversdorf LJ., Chaston SD., Miller TR., McMahon KD. 2015. Microcystin mcyA and mcyE gene abundances are not appropriate indicators of microcystin concentrations in lakes. PLOS ONE 10:1–18. DOI: 10.1371/journal.pone.0125353.

Beversdorf LJ., Miller TR., McMahon KD. 2013. The Role of Nitrogen Fixation in Cyanobacterial Bloom Toxicity in a Temperate, Eutrophic Lake. PLOS ONE 8:1–11. DOI: 10.1371/journal.pone.0056103.

Boden R., Hutt LP., Rae AW. 2017. Reclassification of Thiobacillus aquaesulis (Wood & Kelly, 1995) as Annwoodia aquaesulis gen. nov., comb. nov., transfer of Thiobacillus (Beijerinck, 1904) from the Hydrogenophilales to the Nitrosomonadales, proposal of Hydrogenophilalia class. nov. withi. International Journal of Systematic and Evolutionary Microbiology 67:1191–1205. DOI: 10.1099/ijsem.0.001927.

Boisvert S., Raymond F., Godzaridis É., Laviolette F., Corbeil J. 2012. Ray Meta: scalable de novo metagenome assembly and profiling. Genome Biology 13:1–13. DOI: 10.1186/gb-2012-13-12-r122.

Bowers RM., Kyrpides NC., Stepanauskas R., Harmon-Smith M., Doud D., Reddy TBK., Schulz F., Jarett J., Rivers AR., Eloe-Fadrosh EA., Tringe SG., Ivanova NN., Copeland A., Clum A., Becraft ED., Malmstrom RR., Birren B., Podar M., Bork P., Weinstock GM., Garrity GM., Dodsworth JA., Yooseph S., Sutton G., Glöckner FO., Gilbert JA., Nelson WC., Hallam SJ., Jungbluth SP., Ettema TJG., Tighe S., Konstantinidis KT., Liu WT., Baker BJ., Rattei T., Eisen JA., Hedlund B., McMahon KD., Fierer N., Knight R., Finn R., Cochrane G., Karsch-Mizrachi I., Tyson GW., Rinke C., Lapidus A., Meyer F., Yilmaz P., Parks DH., Eren AM., Schriml L., Banfield JF., Hugenholtz P., Woyke T. 2017. Minimum information about a single amplified genome (MISAG) and a metagenome-assembled genome (MIMAG) of bacteria and archaea. Nature Biotechnology 35:725–731. DOI: 10.1038/nbt.3893.

Bowles MW., Mogollon JM., Kasten S., Zabel M., Hinrichs K-U. 2014. Global rates of marine sulfate reduction and implications for sub-sea-floor metabolic activities. Science Express Reports. DOI: 10.1038/35351.

Bowman JP., Sly LI., Stackebrandt E. 1995. The phylogenetic position of the family Methylococcaceae. International Journal of Systematic Bacteriology 45:182–5. DOI: 10.1099/00207713-45-3-622a.

Bragg JG., Wagner A. 2009. Protein material costs: single atoms can make an evolutionary difference. Trends in Genetics 25:5–8. DOI: 10.1016/j.tig.2008.10.011.

Al Bulushi I., Poole S., Deeth HC., Dykes GA. 2009. Biogenic amines in fish: Roles in intoxication, spoilage, and nitrosamine formation-A review. Critical Reviews in Food Science and Nutrition 49:369–377. DOI: 10.1080/10408390802067514.

Butman D., Stackpoole S., Stets E., McDonald CP., Clow DW., Striegl RG. 2015. Aquatic carbon cycling in the conterminous United States and implications for terrestrial carbon accounting. Proceedings of the National Academy of Sciences: 1–6. DOI: 10.1073/pnas.1512651112.

Camacho C., Coulouris G., Avagyan V., Ma N., Papadopoulos J., Bealer K., Madden TL. 2009. BLAST plus☐: architecture and applications. BMC Bioinformatics 10:1–9. DOI: Artn 421\nDoi 10.1186/1471-2105-10-421.

Caporaso JG., Lauber CL., Walters WA., Berg-Lyons D., Huntley J., Fierer N., Owens SM., Betley J., Fraser L., Bauer M., Gormley N., Gilbert JA., Smith G., Knight R. 2012. Ultra-high-throughput microbial community analysis on the Illumina HiSeq and MiSeq platforms. The ISME Journal 6:1621–1624. DOI: 10.1038/ismej.2012.8.

Chistoserdova L., Kalyuzhnaya MG., Lidstrom ME. 2009. The Expanding World of Methylotrophic Metabolism. Annual Review of Microbiology 63:477–499. DOI: 10.1146/annurev.micro.091208.073600.

Darling AE., Jospin G., Lowe E., Matsen FA., Bik HM., Eisen JA. 2014. PhyloSift: phylogenetic analysis of genomes and metagenomes. PeerJ 2:e243. DOI: 10.7717/peerj.243.

DeSantis TZ., Hugenholtz P., Larsen N., Rojas M., Brodie EL., Keller K., Huber T., Dalevi D., Hu P., Andersen GL. 2006. Greengenes, a chimera-checked 16S rRNA gene database and workbench compatible with ARB. Applied and Environmental Microbiology 72:5069–5072. DOI: 10.1128/AEM.03006-05.

Eiler A., Mondav R., Sinclair L., Fernandez-Vidal L., Scofield D., Scwientek P., Martinez-Garcia M., Torrents D., McMahon KD., Andersson SGE., Stepanauskas R., Woyke T., Bertilsson S. 2016. Tuning fresh: radiation through rewiring of central metabolism in streamlined bacteria. The ISME Journal 10:1–13. DOI: 10.13140/RG.2.1.1968.9040.

Engelbrektson AL., Kunin V., Wrighton KC., Zvenigorodsky N., Chen F., Ochman H., Hugenholtz P. 2010. Experimental factors affecting PCR-based estimates of microbial species richness and evenness. The ISME Journal 4:642.

Ghylin TW., Garcia SL., Moya F., Oyserman BO., Schwientek P., Forest KT., Mutschler J., Dwulit-Smith J., Chan L-K., Martinez-Garcia M., Sczyrba A., Stepanauskas R., Grossart H-P., Woyke T., Warnecke F., Malmstrom R., Bertilsson S., McMahon KD. 2014. Comparative single-cell genomics reveals potential ecological niches for the freshwater acI Actinobacteria lineage. The ISME Journal 8:2503–16. DOI: 10.1038/ismej.2014.135.

del Giorgio PA., Cole JJ., Cimbleris A. 1997. Respiration rates in bacteria exceed phytoplankton production in unproductive aquatic systems. Nature 385:148–151.

Giroldo D., Augusto A., Vieira H. 2005. Polymeric and free sugars released by three phytoplanktonic species from a freshwater tropical eutrophic reservoir. Journal of Plankton Research 27:695–705. DOI: 10.1093/plankt/fbi043.

Gong X., Garcia-Robledo E., Revsbech N-P., Schramm A. 2018. Gene Expression of Terminal Oxidases in Two Marine Bacterial Strains Exposed to Nanomolar Oxygen Concentrations. FEMS Microbiology Ecology. DOI: 10.1093/femsec/fiy072/4983120.

Hall MW., Rohwer RR., Perrie J., Mcmahon KD., Beiko RG. 2017. Ananke☐: temporal clustering reveals ecological dynamics of microbial communities. PeerJ 5:1–19. DOI: 10.7717/peerj.3812.

Hamilton JJ., Garcia SL., Brown BS., Oyserman BO., Moya-Flores F., Bertilsson S., McMahon Katherine D. 2017. Metabolic Network Analysis and Metatranscriptomics Reveal Auxotrophies and Nutrient Sources of the Cosmopolitan Freshwater Microbial Lineage acI. mSystems 2:1–13.

Hanson TE., Tabita FR. 2001. A ribulose-1,5-bisphosphate carboxylase/oxygenase (RubisCO)-like protein from Chlorobium tepidum that is involved with sulfur metabolism and the response to oxidative stress. Proceedings of the National Academy of Sciences 98:4397–4402. DOI: 10.1073/pnas.081610398.

He S., Stevens SL., Chan L-K., Bertilsson S., Glavina Del Rio T., Tringe SG., Malmstrom RR., McMahon KD. 2017. Ecophysiology of Freshwater Verrucomicrobia Inferred from Metagenome-Assembled Genomes. mSphere 2:1–17.

Holkenbrink C., Barbas SO., Mellerup A., Otaki H., Frigaard NU. 2011. Sulfur globule oxidation in green sulfur bacteria is dependent on the dissimilatory sulfite reductase system. Microbiology 157:1229–1239. DOI: 10.1099/mic.0.044669-0.

Hong S., Bunge J., Leslin C., Jeon S., Epstein SS. 2009. Polymerase chain reaction primers miss half of rRNA microbial diversity. The ISME Journal 3:1365–1373. DOI: 10.1038/ismej.2009.89.

Hyatt D., Chen GL., LoCascio PF., Land ML., Larimer FW., Hauser LJ. 2010. Prodigal: Prokaryotic gene recognition and translation initiation site identification. BMC Bioinformatics 11. DOI: 10.1186/1471-2105-11-119.

Igarashi K., Kashiwagi K. 1999. Polyamine transport in bacteria and yeast. Biochem. J. 344:633–642.

Jorgenson NO., Tranvik LJ., Edling H., Graneli W., Lindell M. 1998. Effects of sunlight on occurrence and bacterial turnover of specific carbon and nitrogen compounds in lake water. FEMS Microbiology Ecology 25:217–227.

Kalyuzhnaya MG., Beck DAC., Vorobev A., Smalley N., Kunkel DD., Lidstrom ME., Chistoserdova L. 2011. Novel methylotrophic isolates from lake sediment, description of Methylotenera versatilis sp. nov. and emended description of the genus Methylotenera. International Journal of Systematic and Evolutionary Microbiology 62:106–111. DOI: 10.1099/ijs.0.029165-0.

Kanao T., Kawamura M., Fukui T., Atomi H., Imanaka T. 2002. Characterization of isocitrate dehydrogenase from the green sulfur bacterium Chlorobium limicola: A carbon dioxide-fixing enzyme in the reductive tricarboxylic acid cycle. European Journal of Biochemistry 269:1926–1931. DOI: 10.1046/j.1432-1327.2002.02849.x.

Kang DD., Froula J., Egan R., Wang Z. 2015. MetaBAT, an efficient tool for accurately reconstructing single genomes from complex microbial communities. PeerJ 3:e1165. DOI: 10.7717/peerj.1165.

Karhunen J., Arvola L., Peura S., Tiirola M. 2013. Green sulphur bacteria as a component of the photosynthetic plankton community in small dimictic humic lakes with an anoxic hypolimnion. Aquatic Microbial Ecology 68:267–272. DOI: 10.3354/ame01620.

Latypova E., Yang S., Wang Y., Wang T., Chavkin TA., Hackett M., Schäfer H., Kalyuzhnaya MG. 2010. Genetics of the glutamate-mediated methylamine utilization pathway in the facultative methylotrophic beta-proteobacterium Methyloversatilis universalis FAM5. Molecular Microbiology 75:426–439. DOI: 10.1111/j.1365-2958.2009.06989.x.

Li H., Durbin R. 2010. Fast and accurate long-read alignment with Burrows-Wheeler transform. Bioinformatics 26:589–595. DOI: 10.1093/bioinformatics/btp698.

Linz AM., Crary BC., Shade A., Owens S., Gilbert JA., Knight R., McMahon KD. 2017. Bacterial Community Composition and Dynamics Spanning Five Years in Freshwater Bog Lakes. mSphere 2:1–15. DOI: e00169-17.

Luo R., Liu B., Xie Y., Li Z., Huang W., Yuan J., He G., Chen Y., Pan Q., Liu Y., Tang J., Wu G., Zhang H., Shi Y., Liu Y., Yu C., Wang B., Lu Y., Han C., Cheung DW., Yiu S-M., Peng S., Xiaoqian Z., Liu G., Liao X., Li Y., Yang H., Wang J., Lam T-W., Wang J. 2012. SOAPdenovo2: an empirically improved memory-efficient short-read de novo assembler. GigaScience 1:1–6. DOI: 10.1186/2047-217X-1-18.

Magooc T., Salzberg SL. 2011. FLASH: Fast length adjustment of short reads to improve genome assemblies. Bioinformatics 27:2957–2963. DOI: 10.1093/bioinformatics/btr507.

Markowitz VM., Chen IMA., Palaniappan K., Chu K., Szeto E., Grechkin Y., Ratner A., Jacob B., Huang J., Williams P., Huntemann M., Anderson I., Mavromatis K., Ivanova NN., Kyrpides NC. 2012. IMG: The Integrated Microbial Genomes database and comparative analysis system. Nucleic Acids Research 40:115–122. DOI: 10.1093/nar/gkr1044.

Martinez-Garcia M., Swan BK., Poulton NJ., Gomez ML., Masland D., Sieracki ME., Stepanauskas R. 2012. High-throughput single-cell sequencing identifies photoheterotrophs and chemoautotrophs in freshwater bacterioplankton. The ISME Journal 6:113–123. DOI: 10.1038/ismej.2011.84.

Mou X., Vila-Costa M., Sun S., Zhao W., Sharma S., Moran MA. 2011. Metatranscriptomic signature of exogenous polyamine utilization by coastal bacterioplankton. Environmental Microbiology 3:798–806. DOI: 10.1111/j.1758-2229.2011.00289.x.

Newton RJ., Jones SE., Eiler A., McMahon KD., Bertilsson S. 2011. A guide to the natural history of freshwater lake bacteria. Microbiology and Molecular Biology Reviews 75:14–49. DOI: 10.1128/MMBR.00028-10.

Parks DH., Imelfort M., Skennerton CT., Hugenholtz P., Tyson GW. 2015. CheckM: assessing the quality of microbial genomes recovered from isolates, single cells, and metagenomes. Genome Research 25:1043–1055.

Paver SF., Kent AD. 2017. Temporal Patterns in Glycolate-Utilizing Bacterial Community Composition Correlate with Phytoplankton Population Dynamics in Humic Lakes. Microbial Ecology 60:406–418. DOI: 10.1007/S00248-0.

Peters JW., Schut GJ., Boyd ES., Mulder DW., Shepard EM., Broderick JB., King PW., Adams MWW. 2015. [FeFe]-and [NiFe]-hydrogenase diversity, mechanism, and maturation. Biochimica et Biophysica Acta - Molecular Cell Research 1853:1350–1369. DOI: 10.1016/j.bbamcr.2014.11.021.

Peura S., Eiler A., Bertilsson S., Nyka H., Tiirola M., Jones RI. 2012. Distinct and diverse anaerobic bacterial communities in boreal lakes dominated by candidate division OD1. The ISME Journal 6:1640–1652. DOI: 10.1038/ismej.2012.21.

Peura S., Sinclair L., Bertilsson S., Eiler A. 2015. Metagenomic insights into strategies of aerobic and anaerobic carbon and nitrogen transformation in boreal lakes. Scientific Reports 5:12102. DOI: 10.1038/srep12102.

Pomeroy LR., Wiebe WJ. 1988. Energetics of microbial food webs. Hydrobiologia 159:7–18. DOI: 10.1007/BF00007363.

Price MN., Dehal PS., Arkin AP. 2010. FastTree 2 - Approximately maximum-likelihood trees for large alignments. PLOS ONE 5. DOI: 10.1371/journal.pone.0009490.

Ramachandran A., Walsh DA. 2015. Investigation of XoxF methanol dehydrogenases reveals new methylotrophic bacteria in pelagic marine and freshwater ecosystems. FEMS Microbiology Ecology 91. DOI: 10.1093/femsec/fiv105.

Ramanan R., Kim B-H., Cho D-H., Oh H-M., Kim H-S. 2015. Algae-bacteria interactions: evolution, ecology and emerging applications. Biotechnology Advances. DOI: 10.1016/j.biotechadv.2015.12.003.

Remsen CC., Carpenter EJ., Schroeder BW. 1972. Competition for Urea among Estuarine Microorganisms. Ecological Society of America 53:921–926.

Rinke C., Schwientek P., Sczyrba A., Ivanova NN., Anderson IJ., Cheng J-F., Darling AE., Malfatti S., Swan BK., Gies E a., Dodsworth J a., Hedlund BP., Tsiamis G., Sievert SM., Liu W-T., Eisen J a., Hallam SJ., Kyrpides NC., Stepanauskas R., Rubin EM., Hugenholtz P., Woyke T. 2013. Insights into the phylogeny and coding potential of microbial dark matter. Nature 499:431–437. DOI: 10.1038/nature12352.

Rognes T., Flouri T., Nichols B., Quince C., Mahé F. 2016. VSEARCH: a versatile open source tool for metagenomics. PeerJ 4:1–18. DOI: 10.7717/peerj.2584.

Rohwer RR., Hamilton JJ., Newton RJ., McMahon KD. 2017. TaxAss☐: Leveraging a Custom Freshwater Achieves Fine-Scale Taxonomic Resolution. bioRxiv.

Roux S., Chan LK., Egan R., Malmstrom RR., McMahon KD., Sullivan MB. 2017. Ecogenomics of virophages and their giant virus hosts assessed through time series metagenomics. Nature Communications 8. DOI: 10.1038/s41467-017-01086-2.

Salcher MM., Neuenschwander SM., Posch T., Pernthaler J. 2015. The ecology of pelagic freshwater methylotrophs assessed by a high-resolution monitoring and isolation campaign. The ISME Journal 9:2442–2453. DOI: 10.1038/ismej.2015.55.

Salcher MM., Posch T., Pernthaler J. 2013. In situ substrate preferences of abundant bacterioplankton populations in a prealpine freshwater lake. Isme J 7:896–907. DOI: 10.1038/ismej.2012.162.

Schloss PD., Westcott SL., Ryabin T., Hall JR., Hartmann M., Hollister EB., Lesniewski RA., Oakley BB., Parks DH., Robinson CJ., Sahl JW., Stres B., Thallinger GG., Van Horn DJ., Weber CF. 2009. Introducing mothur: Open-source, platform-independent, community-supported software for describing and comparing microbial communities. Applied and Environmental Microbiology 75:7537–7541. DOI: 10.1128/AEM.01541-09.

Segata N., Waldron L., Ballarini A., Narasimhan V., Jousson O., Huttenhower C. 2012. Metagenomic microbial community profiling using unique clade-specific marker genes. Nature Methods 9:811–4. DOI: 10.1038/nmeth.2066.

Šimek K., Kasalický V., Jezbera J., Jezberová J., Hejzlar J., Hahn MW. 2010. Broad habitat range of the phylogenetically narrow R-BT065 cluster, representing a core group of the betaproteobacterial genus limnohabitans. Applied and Environmental Microbiology 76:631–639. DOI: 10.1128/AEM.02203-09.

Sommer DD., Delcher AL., Salzberg SL., Pop M. 2007. Minimus: a fast, lightweight genome assembler. BMC Bioinformatics 8:64. DOI: 10.1186/1471-2105-8-64.

Tang KH., Blankenship RE. 2010. Both forward and reverse TCA cycles operate in green sulfur bacteria. Journal of Biological Chemistry 285:35848–35854. DOI: 10.1074/jbc.M110.157834.

Varghese NJ., Mukherjee S., Ivanova N., Konstantinidis KT., Mavrommatis K., Kyrpides NC., Pati A. 2015. Microbial species delineation using whole genome sequences. Nucleic Acids Research 43:gkv657-. DOI: 10.1093/nar/gkv657.

Walsh JR., Munoz SE., Vander Zanden MJ. 2016. Outbreak of an undetected invasive species triggered by a climate anomaly. Ecosphere 7:1–17. DOI: 10.1002/ecs2.1628.

Williamson CE., Dodds W., Kratz TK., Palmer MA. 2008. Lakes and streams as sentinels of environmental change in terrestrial and atmospheric processes. Frontiers in Ecology and the Environment 6:247–254. DOI: 10.1890/070140.

Yin Y., Mao X., Yang J., Chen X., Mao F., Xu Y. 2012. DbCAN: A web resource for automated carbohydrate-active enzyme annotation. Nucleic Acids Research 40:445–451. DOI: 10.1093/nar/gks479.

